# Cortical plasticity of the tactile mirror system in borderline personality disorder

**DOI:** 10.64898/2026.01.16.699954

**Authors:** Agnese Zazio, Giacomo Guidali, Cora Miranda Lanza, Elisa Dognini, Christian Mancini, Serena Meloni, Barbara Borroni, Roberta Rossi, Nadia Bolognini, Marta Bortoletto

## Abstract

Individuals with borderline personality disorder (BPD) show alterations in empathic abilities, potentially involving automatic simulation processes supported by mirror-like mechanisms in the somatosensory domain. Within the tactile mirror system (TaMS), observing touch on another person’s body activates cortical regions involved in tactile perception, including the primary somatosensory cortex (S1). Although mirror-like alterations have been suggested in BPD, the underlying mechanisms of plasticity remain underexplored. Here, we used a cross-modal paired associative stimulation (cm-PAS) protocol to investigate the plasticity mechanisms of TaMS functioning in BPD. Twenty-four individuals with BPD and 24 healthy controls (HCs) were included. Empathic abilities were assessed using self-report questionnaires. Participants performed tactile acuity and visuo-tactile spatial congruity (VTSC) tasks before and after a cm-PAS protocol. During cm-PAS, images of a hand being touched were paired with transcranial magnetic stimulation over the S1. The effects of cm-PAS were assessed on tactile acuity, as an index of S1 activity, and VTSC performance, as an index of TaMS functioning. Preregistered analyses revealed that patients with BPD tended to have lower cognitive empathy than HCs, with no significant cm-PAS effects on tactile acuity or VTSC performance in HCs, precluding between-group comparisons of plasticity effects. Exploratory analyses were conducted to further investigate potential sources of variability in the effects of cm-PAS, as well as the relationship between cognitive empathy and visuo-tactile processing as measure of TaMS functioning. Preregistered Stage 1 protocol: https://osf.io/sqnwd (date of in-principle acceptance: 25/06/2023).

## Introduction

Borderline personality disorder (BPD) is a mental health problem characterized by a pervasive pattern of impulsivity, self-image issues, difficulties in emotion regulation and unstable interpersonal relationships (Lazarus, Cheavens, Festa, & Zachary Rosenthal, 2014). Interpersonal dysfunctions are one of the main features of the disorder and have been associated with difficulties in mentalization as well as in the ability to empathize with others, with greater deficits in the cognitive dimension (i.e., understanding others’ perspective) rather than in the affective dimension (sensing others’ feelings; (Harari, Shamay-Tsoory, Ravid, & Levkovitz, 2010; F. Martin, Flasbeck, Brown, & Brüne, 2017). It has been suggested that empathic abilities may involve automatic simulation processes relying on mirror-like mechanisms, not only in the motor domain but also in the somatosensory domain (Bolognini, Rossetti, Convento, & Vallar, 2013; Keysers & Gazzola, 2009; Keysers, Kaas, & Gazzola, 2010). In the so-called tactile mirror system (TaMS), the observation of a touch on someone else’s body has been shown to activate the same cortical network involved in tactile perception, including the primary somatosensory cortex (S1; (Bolognini, Rossetti, Fusaro, Vallar, & Miniussi, 2014; Pisoni, Romero Lauro, Vergallito, Maddaluno, & Bolognini, 2018). Intriguingly, the activity of TaMS has been shown to correlate with measures of cognitive empathy in the healthy population (Bolognini, Miniussi, Gallo, & Vallar, 2013; Bolognini et al., 2014), thus suggesting the hypothesis that the difficulties in cognitive empathy observed in BPD patients may be associated with alterations in TaMS functioning.

While previous studies have suggested mirror system alterations in BPD (Mier et al., 2013; Sosic-Vasic et al., 2019), still little is known about the integrity of a basic neurophysiological mechanism such as brain plasticity within these systems. Considering that the development of mentalization and empathic abilities appears to rely on early associative learning and metaplasticity, it has been suggested that the pathophysiology of BPD may be associated with alterations in neuronal plasticity, mediated by N-methyl-D-aspartate (NMDA) neurotransmission; however, evidence is still lacking (Grosjean & Tsai, 2007). Importantly, the integrity of plasticity mechanisms may represent a critical factor for the effectiveness of therapeutic interventions (Mancke et al., 2018).

Paired associative stimulation (PAS) protocols represent a well-established tool to non-invasively induce brain plasticity effects in humans (Guidali, Roncoroni, & Bolognini, 2021; Stefan, Kunesch, Cohen, Benecke, & Classen, 2000; Suppa et al., 2017; Wischnewski & Schutter, 2016). To induce changes in synaptic efficacy, PAS protocols exploit the Hebbian learning rule and the concept of spike-timing dependent plasticity observed at the cellular level, namely that neural connections are strengthened (or weakened) in the case of repeated activations of the presynaptic neuron before the postsynaptic neuron (or vice-versa) in a critical time interval of a few tens of ms (Feldman, 2012; Hebb, 1949; Miller, 1996). In classical PAS protocols targeting the somatosensory system (S1-PAS), a peripheral electrical stimulus over the wrist (acting as presynaptic activation) is repeatedly paired with a transcranial magnetic stimulation (TMS) pulse over S1 in the contralateral hemisphere (acting as postsynaptic activation). Depending on the time interval between the two stimuli, the S1-PAS may induce long-term potentiation or depression (LTP- or LTD-like, respectively) effects, lasting up to 30 minutes after protocol delivery (Alexander Wolters et al., 2005). In addition to neurophysiological effects (i.e., somatosensory evoked potential modulation), these plastic mechanisms have also been detected by exploiting behavioral measures of S1 functioning, such as tactile acuity (Litvak et al., 2007).

PAS protocols have also been used to target cross-modal networks, since they pairs stimulations, the peripheral and cortical stimuli, belonging to different systems (e.g., Guidali, Carneiro, & Bolognini, 2020; Sowman, Dueholm, Rasmussen, & Mrachacz-Kersting, 2014; Suppa, Li Voti, Rocchi, Papazachariadis, & Berardelli, 2015). Recently, a cross-modal visuo-tactile PAS (cm-PAS) has been developed with the aim of targeting the TaMS (Maddaluno, Guidali, Zazio, Miniussi, & Bolognini, 2020; Zazio, Guidali, Maddaluno, Miniussi, & Bolognini, 2019). Compared to classical S1-PAS, in cm-PAS the peripheral electrical stimulus on the wrist is replaced by a visual stimulus of a hand being touched. The efficacy of the cm-PAS in inducing LTP-like mechanisms has been shown in a series of experiments, consisting of an increase in tactile acuity that was specific for the time interval between the visual stimulus and the TMS (i.e., 20 ms and not 60 nor 100 ms), for the site of cortical stimulation (i.e., S1 and not the primary visual cortex), and for the content of the visual stimulus (i.e., a hand being touched and not a moving hand). Moreover, cm-PAS modulated a neurophysiological correlate of S1 activity, namely, the amplitude of the P40 component of somatosensory-evoked potentials increased after cm-PAS. Overall, these findings are consistent with the hypothesis that when seeing a human touch on someone else’s body, S1 is recruited by mirror-like mechanisms and can be involved in plasticity mechanisms (Zazio et al., 2019).

Taking together the existing evidence, here, we aim to shed light on the neural basis of interpersonal dysfunction in BPD by bridging the gap between the literature on empathic alterations in BPD patients, on the one hand, and on the neurophysiological underpinnings of TaMS and its plastic properties in the healthy population, on the other hand. Specifically, we investigate the integrity of plastic modulations within the TaMS in BPD patients, by employing the previously described cm-PAS.

The present study involved BPD patients and healthy controls (HCs) undergoing two sessions of cm-PAS, i.e., an experimental session and a control session with a different time interval between the paired stimuli. The two groups have been compared in the cognitive dimensions of empathic abilities, measured by means of a self-report questionnaire. To assess the effects of the cm-PAS, both groups underwent a two-point discrimination task (2-PDT) as a measure of tactile acuity, an index of S1 activity (Litvak et al., 2007), before and after the cm-PAS protocol. Moreover, to explore whether the effects of the cm-PAS are limited to S1 activity or extend to behavioral correlates of TaMS, both groups also performed a visuo-tactile spatial congruity (VTSC) task before and after the cm-PAS protocol, to measure TaMS functioning (Bolognini et al., 2014).

The hypotheses of the present study are the following (see Study design template in **Table 1**):

I. BPD patients are expected to show reduced cognitive empathy compared to HCs, as measured by means of a self-report questionnaire, consistently with previous findings (Grzegorzewski, Kulesza, Pluta, Iqbal, & Kucharska, 2019; Harari et al., 2010; F. Martin et al., 2017).
II. HCs are expected to show an improvement in tactile acuity after cm-PAS. This effect is considered a positive control, as it supports the effectiveness of a PAS protocol relying on TaMS (i.e., cm-PAS) in inducing plastic mechanisms (Zazio et al., 2019).
III. The effects of cm-PAS are expected to extend to a behavioral measure of TaMS functioning, i.e., the VTSC task: the cm-PAS is expected to strengthen the association between visual touches and S1 activation, thus inducing a decrease in performance, arising from a greater interference in case of spatial incongruity between the visual and the real touches and/or a greater facilitation in congruent trials. Therefore, we hypothesize HCs to show a decrease in performance on VTSC task after cm-PAS, as indexed by a greater difference in reaction times (RTs) between incongruent and congruent trials. No effects are expected after the control session of cm-PAS.
IV. In BPD, we hypothesize an alteration of plasticity within TaMS. Considering the empathic alterations in BPD, the rationale is the following. On the one hand, the development of mentalization and empathic abilities appears to rely on early associative learning (Grosjean & Tsai, 2007), on the other hand, mirror system alterations associated with empathic dysfunctions have been reported in BPD; therefore, we hypothesize that the pathophysiology of BPD may be associated with alterations in neuronal plasticity within TaMS (Mier et al., 2013; Sosic-Vasic et al., 2019). Specifically, we expect a reduced or null effect of cm-PAS compared to HCs on tactile acuity and on VTSC task. No effects are expected after the control session of cm-PAS.

**Table 1.** Study Design Template. Abbreviations list: BPD = Borderline Personality Disorder; cm-PAS = Cross-Modal Paired Associative Stimulation; HC = Healthy Control; ISI = Inter Stimulus Interval; LTP = Long-Term Potentiation; QCAE = Questionnaire of Cognitive and Affective Empathy; TaMS = Tactile Mirror System; TMS = Transcranial Magnetic Stimulation; VTSC = Visuo-Tactile Spatial Congruity.

| Question | Hypothesis | Sampling plan | Analysis Plan | Rationale for deciding the sensitivity of the test for confirming or disconfirming the hypothesis | Interpretation given different outcomes | Observed outcome |
| --- | --- | --- | --- | --- | --- | --- |
| *I. Do patients with BPD show alterations in their empathic abilities?* | Cognitive empathy: BPD < HCs.<br>Patients with BPD will show lower cognitive empathy levels (i.e., mean score given by the subscales 'perspective taking' and 'online simulation') compared to HCs, as assessed by means of QCAE. | Power analysis based on preliminary data collected by the authors for another study (not published yet at the time of the Stage 1 submission; Zazio et al., 2024). Cognitive empathy measured with the QCAE has been compared between 18 BPD patients and 15 HCs. Power analysis has been performed in GPower (v. 3.1.9.7, Faul et al. 2007), with a power of 90% and a level of statistical significance of 2%.<br>BPD patients show lower cognitive empathy compared to HCs ($d = 1.07$). The estimated sample for a one-tailed independent t-test resulted in 21 participants per group. | 24 patients with BPD and 24 HCs will be compared in cognitive empathy levels obtained from the QCAE by means of a one-tailed Yuen's trimmed mean t-test, a robust version of the t-test for independent samples (Mair & Wilcox, 2020). | The effect size for power analysis was determined based on preliminary data collected at the IRCCS Fatebenefratelli in Brescia (where the experiment will take place), as the BPD group is likely to be representative of the patients that will be tested in the present study. | Lower cognitive empathy levels in self-report measures in patients with BPD compared to HCs will be interpreted as a difficulty in understanding others' mental states. If the two groups will not differ in the empathic abilities, it will be discussed whether the obtained results represent a non-replication of previous findings or a difference in the samples. | Unsupported. The comparison between patients with BPD and HCs on QCAE cognitive empathy scores showed a trend toward significance in the expected direction (HCs > patients with BPD; $t = 1.95$, $p = 0.031$). |
| *II. Is cm-PAS<sub>ISI-20</sub> effective in inducing plastic mechanisms in HCs, as indexed by tactile acuity?*<br>(positive control) | Improvement in tactile acuity after cm-PAS in HCs: cm-PAS<sub>ISI-20</sub> > cm-PAS<sub>ISI-100</sub>.<br>HCs will show a greater improvement in tactile acuity (i.e., $d'$ sensory threshold) after the cm-PAS<sub>ISI-20</sub> compared to the control condition cm-PAS<sub>ISI-100</sub>. | Power analysis based on previous findings on the effects of cm-PAS in HCs on tactile acuity (Zazio et al., 2019), performed in GPower (v. 3.1.9.7, Faul et al. 2007), with a power of 90% and a level of statistical significance of 2%.<br>We considered the data collected in Experiment 1, which included the same cm-PAS conditions as in the present study. To estimate the power needed to detect the interaction effect of within-subject factors, we considered the difference in sensory threshold ($d'$ values), i.e., $\Delta\text{Post}_{\text{cm-PAS}} - \text{Pre}_{\text{cm-PAS}}$, in the two PAS conditions (effect size $dz = 0.95$) and tested power for a one-tailed paired t-test. The estimated sample resulted in 15 participants. | A rm-ANOVA with factors Time ($\text{Pre}_{\text{cm-PAS}}$, $\text{Post}_{\text{cm-PAS}}$) and ISI (ISI-20, ISI-100) will be performed on tactile acuity recorded in 24 HCs. | The effect size for power analysis was determined based on a previous study by Zazio et al. (2019), testing the effects of cm-PAS on tactile acuity in HCs, which included the same cm-PAS paradigm as well as the same control condition of the present study. | An interaction Time X ISI showing a greater improvement in tactile acuity after cm-PAS<sub>ISI-20</sub> compared to cm-PAS<sub>ISI-100</sub> will be interpreted as a successful replication of previous findings, supporting the efficacy of cm-PAS in inducing LTP-like mechanisms within the somatosensory domain.<br>The absence of a Time X ISI interaction will not confirm the hypothesis, and hypothesis IV will not be tested. | Unsupported. No significant Time × ISI interaction was observed ($p = 0.921$), failing to replicate previous cm-PAS effects on tactile acuity. |
| *III. Is cm-PAS<sub>ISI-20</sub> effective in inducing plastic mechanisms in HCs, as indexed by behavioral performance on VTSC task?* | Decrease in performance (i.e., greater $\Delta\text{RT}_{\text{incong-cong}}$) on VTSC task in HC after cm-PAS: cm-PAS<sub>ISI-20</sub> > after cm-PAS<sub>ISI-100</sub>.<br>HCs will show a greater decrease in VTSC task performance | Power analysis considered the same parameters described in hypothesis II. | A rm-ANOVA with factors Time ($\text{Pre}_{\text{cm-PAS}}$, $\text{Post}_{\text{cm-PAS}}$) and ISI (ISI-20, ISI-100) will be performed on VTSC measures recorded in 24 HCs. | The effect size for power analysis was determined based on previous findings of cm-PAS on tactile acuity, as there are no studies testing the efficacy of cm-PAS on VTSC. Nevertheless, a previous study using a similar VTSC task and using an online TMS protocol to induce modulations of performance (Bolognini et al., 2014), observed | An interaction Time X ISI showing stronger decrease in behavioral performance after cm-PAS<sub>ISI-20</sub> than after PAS<sub>ISI-100</sub> will support the efficacy of cm-PAS in inducing LTP-like plasticity mechanisms that extends to behavioral measures of TaMS functioning.<br>The absence of a Time X ISI interaction will not confirm the hypothesis, | Unsupported. No significant Time × ISI interaction was observed ($p = 0.328$), indicating no cm-PAS effects on behavioral measures of TaMS functioning as indexed |
| | after the cm-PAS<sub>ISI-20</sub> compared to the control condition cm-PAS<sub>ISI-100</sub>. | | | significant effects on RTs with an effect size ranging between $\eta p^2 = 0.25-0.31$ for a rm-ANOVA interaction, which is similar to the one observed for tactile acuity in Zazio et al. 2019 ($\eta p^2 = 0.3$). | and hypothesis IV will not be tested. | by VTSC task. |
| *IV. Are plasticity mechanisms within the TaMS reduced in BPD patients?* | Improvement in tactile acuity after cm-PAS: BPD < HCs.<br>Decrease in VTSC task performance after cm-PAS: BPD < HCs.<br>Compared to HCs, BPD patients will show reduced or null effect of cm-PAS, both on tactile acuity (i.e., $d'$ sensory threshold) and performance on the VTSC task (i.e., $\Delta RT_{incong-cong}$). | Power analysis performed in GPower (v. 3.1.9.7, Faul et al. 2007) considered a medium effect size ($\eta p^2 = 0.06$), with a power of 90% and a level of statistical significance of 2%. Based on previous data on HC (Zazio et al. 2019), we considered a correlation between repeated measures of 0.7.<br>The estimated sample size for a between-within interaction rm-ANOVA resulted in 22 participants per group. | 24 HCs and 24 BPD patients will be compared in the effects induced by the cm-PAS on tactile acuity and on the VTSC performance, in separate analyses. The analyses will consist of 2x2x2 mixed-design analysis of variance (ANOVA) with between factor Group (BPD, HCs) and within factors Time (Pre<sub>cm-PAS</sub>, Post<sub>cm-PAS</sub>) and ISI (ISI-20, ISI-100). | In the absence of previous studies testing plasticity within TaMS in BPD, we hypothesized a medium effect size ($\eta p^2 = 0.06$). | An interaction Group X Time X ISI, showing differences between BPD patients and HCs in the effects of cm-PAS, will be interpreted as an alteration of the plastic mechanisms within the TaMS.<br>The absence of an interaction with factor Group will not confirm the hypothesis. | This hypothesis could not be tested due to non significant findings in previous hypotheses II and III. |

We highlight that the present study aims at detecting strong effects, while possible smaller but interesting effects may be missed (see *Sample size estimation*).

## Materials and Methods

### Participants

A total of 36 individuals with BPD and 35 HCs participated in this study (see “Exclusion criteria paragraph” and Figure 1). In the HC group, 10 participants did not complete the experiment because of the exclusion criteria (one for technical issues during the second session, six for stimulation intensity exceeding 90% of the maximal stimulator output, two because of low performance in VTSC catch trials, and one who withdrew after the first session). Among the 25 HCs who completed the experiment, one was excluded because of an outlier performance in the 2-PDT task. In the BPD group, 11 participants did not complete the experiment owing to the exclusion criteria (seven for stimulation intensity exceeding 90% of the maximal stimulator output and four who withdrew after the first session). Among the 25 patients with BPD who completed the experiment, one was excluded as an outlier in the 2-PDT task.

**Figure 1.**
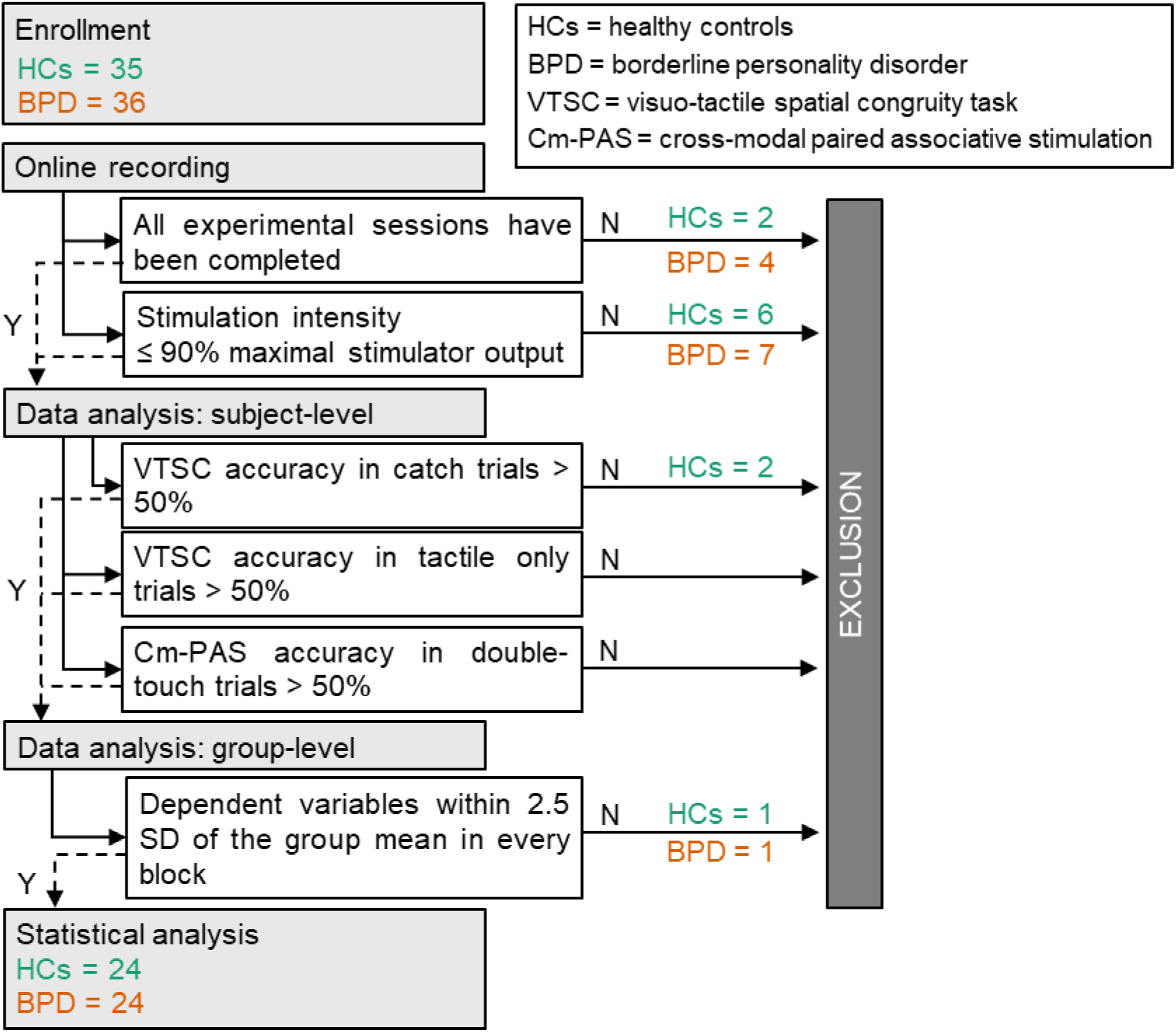
Flowchart of the experimental steps and the corresponding decisions (Y=Yes, N=No) for the inclusion of participants. The number of initially enrolled participants, those excluded, and the final sample are reported. A complete list of study-specific acronyms is provided in the upper-right panel.

As planned, the final sample consisted of 24 individuals with BPD (mean age ± SD: 23.4 years ± 3.2; 21/24 women) matched one-to-one for age and gender with 24 HCs (mean age ± SD: 23.5 ± 4.4; 21/24 women).

Participants were mostly from the local community: BPD patients were recruited by the Unit of Psychiatry, and HCs by word of mouth from the general local population. BPD patients were matched one-to-one with HCs for gender and age (with a tolerability of ± 2 years).

Pharmacological treatments did not represent an exclusion criterion per se, according to the latest TMS guidelines (Rossi et al., 2021). Nevertheless, participants were not included in the study in case they were taking pharmacological treatments that decrease the epileptic threshold (e.g., clozapine, bupropion). Up to now, in the previous experience of the Research Unit of Psychiatry, only 3 out of 200 BPD (1.5%) patients were taking one of these two treatments; therefore, the sample included in the study is representative of the BPD population.

All participants were between 18 and 40 years old, right-handed according to the Edinburgh Handedness Inventory (Oldfield, 1971), with no contraindication to TMS (Rossi et al., 2021). The study was conducted at the Neurophysiology Lab of the IRCCS Istituto Centro San Giovanni di Dio Fatebenefratelli (Brescia, Italy). It has received approval from the local ethics committee (reference number: Parere 65-2020), and it was performed in accordance with the ethical standards of the Declaration of Helsinki.

### Exclusion criteria

Participants were excluded in the following cases: they did not complete experimental sessions; the stimulation intensity in any session exceeded 90% of the maximal stimulator output; accuracy in catch trials during the cm-PAS in any session was below 50%; accuracy during the VTSC in tactile-only trials and/or catch trials was below 50%. Finally, at the group level, participants were excluded in case in any block the dependent variables (see following paragraphs for further details) exceeded ± 2.5 standard deviations (SD) of the group mean.

### Sample size estimation

The sample size estimation was performed for each hypothesis, and we considered the one resulting in the highest sample. In GPower (v. 3.1.9.7, (Faul, Erdfeldfer, Lang, & Buchner, 2007), we set a power of 90% and a level of statistical significance of 2%. Please note that considering the exact effect sizes previously reported, smaller but still meaningful effects may not be detected (Dienes, 2021; Perugini, Gallucci, & Costantini, 2014).

I. For the comparison between BPD and HCs in empathic abilities, the power analysis was based on preliminary data for another study collected by the authors at the IRCCS Fatebenefratelli in Brescia, where the experiment took place, as the BPD group was likely to be representative of the patients that were tested in the present study (not published yet; see pre-registration on Open Science Framework <u>here</u>; by the time of Stage 2, the paper has been published in Zazio et al., 2024. Cognitive empathy measured with the Questionnaire for Cognitive and Affective Empathy (QCAE) has been compared between 18 BPD patients and 15 HCs. BPD patients showed lower cognitive empathy compared to HCs (*d* = 1.07). The estimated sample for the one-tailed independent t-test was 21 participants per group.
II. Regarding the effects of cm-PAS on tactile acuity in healthy participants, we focused on the data collected in a previous study (Zazio et al., 2019; Experiment 1), which included the same cm-PAS conditions as in the present study, namely cm-PAS with an inter-stimulus interval (ISI) of 20 ms, and a control condition of cm-PAS with an ISI of 100 ms. To estimate the power needed to detect the interaction effect of within-subject factors, we considered the difference in sensory threshold (*d’* values), i.e., ΔPost_cm-PAS_-Pre_cm-PAS_, in the two PAS conditions (*dz* = 0.95) and tested power for a one-tailed paired t-test. The estimated sample comprised of 15 participants.
III. For the effects of cm-PAS on VTSC task performance, power analysis considered the same parameters described in hypothesis II. As there are no studies testing the efficacy of cm-PAS on VTSC, the effect size was determined based on previous findings of cm-PAS on tactile acuity. The effect size described for the effect of cm-PAS on tactile acuity (*ηp*^2^ = 0.3; (Zazio et al., 2019) is similar to the one described in a previous study aiming at modulating VTSC performance, as indexed by RTs, by means of online TMS protocols (*ηp*^2^ = 0.25 -0.31; (Bolognini et al., 2014).,
IV. In the absence of previous studies testing plasticity within TaMS in BPD, to assess plasticity alterations we hypothesized a medium effect size (*ηp^2^* = 0.06). Based on previous data on HCs (Zazio et al. 2019), we considered a correlation between repeated measures of 0.7. The estimated sample size for a between-within interaction rm-ANOVA resulted in 22 participants per group. While the use of *a priori* defined levels of effect size may be criticized, the one used here is much smaller than the ones observed in the field (see hypothesis III). While we cannot fully rule out the possibility that smaller effects may be missed, possible negative findings may indicate that alterations of the tactile mirror system are not that crucial for BPD.

Taken together the results of the sample size estimation, the final sample was planned to include 24 participants per group, to ensure a counterbalance of session and task order.

### Clinical assessment

Patient recruitment and clinical assessment were performed by the Research Unit of Psychiatry of the IRCCS Fatebenefratelli. Patients were evaluated using the Structural Clinical Interview for DSM-5 (SCID-CV and SCID-PD) (First, Williams, Karg, & Spitzer, 2017) for the diagnosis of BPD. Patients were not included in case of comorbidity with schizophrenia and other psychotic disorders, according to DSM-5, and in case of unstable pharmacological therapy.

The severity of the symptoms was assessed by means of the Zanarini rating scale for BPD (ZAN-BPD, (Zanarini, 2003)) and the Symptoms Check-list 90 Revised (SCL-90-R, (Derogatis, 1994)). Depressive symptoms were evaluated with the Beck Depression Inventory II (BDI-II, (Beck, 1988)), impulsiveness with the Barratt Impulsiveness Scale (BIS, (Patton, Stanford, & Barratt, 1995)), and alexithymia with the Toronto Alexithymia Scale (TAS-20, (Bagby, Taylor, & Parker, 1994)). Moreover, interpersonal functioning was evaluated with the Inventory of Interpersonal Problems (IIP, (Pilkonis, Kim, Proietti, & Barkham, 1996)), and attachment style was assessed with the Attachment Style Questionnaire (ASQ; (Feeney, Noller, & Hanrahan, 1994)). Finally, the Childhood Trauma Questionnaire (CTQ;(Bernstein & Fink, 1998)) was administered for the assessment of traumatic experiences, and the Inventory of statements about self-injury (ISAS, (Klonsky & Glenn, 2009)) for the evaluation of self-harm. The outcome of scales and questionnaires is reported in aggregated form for descriptive purposes, and possible missing data did not represent an exclusion criterion.

### Design and procedure

Participants were involved in a two-session experiment, on separate days with at least 48 hours between them, at the same time of the day (i.e., in the morning or in the afternoon). Participants were evaluated on their empathic abilities by means of a self-report questionnaire (see below) at the beginning of the first session. Then, the two sessions differed only in the time interval for TMS delivery during the cm-PAS. In both sessions, before and after the cm-PAS protocol, participants underwent a 2-PDT to test for tactile acuity and a VTSC task to test for tactile mirror system functioning, in counterbalanced order; within each participant, the same order was kept before and after the cm-PAS and between sessions (**Figure 2**).

**Figure 2.**
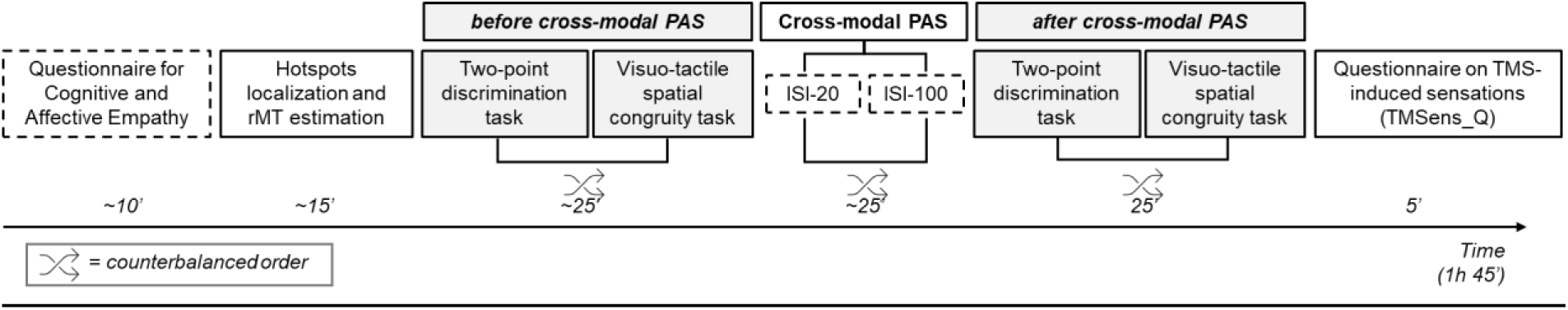
Schematic representation of the experimental session. First, participants filled out the Questionnaire for Cognitive and Affective Empathy on their empathic abilities. Second, the motor hotspot was localized, the resting motor threshold (rMT) was estimated, and then the primary somatosensory cortex (S1) hotspot was identified. Then, baseline tactile acuity at the two-point discrimination task and performance at the tactile acuity and visuo-tactile spatial congruity tasks were recorded (in counterbalanced order), followed by the cross-modal PAS (cm-PAS) with either an ISI of 20 or 100 ms (in separate sessions in counterbalanced order). The two-point discrimination task and the visuo-tactile spatial congruity task were recorded again within 30 minutes after the end of the cm-PAS, following the same order as in the baseline. Finally, participants filled out a self-report questionnaire on sensations induced by TMS (i.e., the TMS_SensQ). Dashed panels: steps performed in one session only; continuous panels: steps performed in each session.

### Questionnaire for empathic abilities

Participants filled out a self-report questionnaire to assess their empathic abilities, namely, the Questionnaire for Cognitive and Affective Empathy (QCAE; (Di Girolamo, Giromini, Winters, Serie, & de Ruiter, 2019; Reniers, Corcoran, Drake, Shryane, & Völlm, 2011)). The QCAE has already been administered in BPD patients (Grzegorzewski et al., 2019), and it has been proposed to overcome some intrinsic limitations of the Interpersonal Reactivity Index (Davis, 1983), another questionnaire on empathic abilities, both from psychometric (Chrysikou & Thompson, 2016) and theoretical (Michaels et al., 2014) points of view. As a dependent variable, we have considered the mean score given by the subscales ‘perspective taking’ and ‘online simulation’.

### Transcranial magnetic stimulation (TMS)

TMS was delivered using a figure-of-eight coil (Magstim model Alpha B.I. Coil Range, diameter: 70 mm) and a monophasic Magstim 2002 stimulator (Magstim, Whitland, UK). First, the motor hotspot for the abductor pollicis brevis (APB) muscle of the left hand was identified in the right hemisphere as the highest and most reliable motor-evoked potentials with the same TMS intensity. Second, the resting motor threshold (rMT) was estimated using the maximum-likelihood threshold hunting algorithm (Awiszus, 2003, 2011), a variant of the best parameter estimation by sequential testing (best PEST procedure; (Pentland, 1980). This procedure has been performed for each participant at the beginning of each session. During this phase, electromyography was visualized online by means of a bipolar belly-tendon montage of the APB of the left hand (g.HIamp, g.tec medical engineering GmbH, Schiedlberg, Austria). Then, right S1 has been localized 2 cm lateral and 0.5 cm posterior to the APB motor hotspot, according to (Holmes & Tamè, 2019; Holmes et al., 2019). During the stimulation of S1, coil orientation was kept approximately at 45° from the midline, inducing a posterior-anterior current direction in the brain. Coil position was monitored online using a neuronavigation system (Softaxic Optic 3.4; EMS, Bologna, Italy). Coil locations were not saved. At the end of the session, participants were asked to fill out a self-report questionnaire on the sensations induced by TMS (TMSens_Q, Giustiniani et al., 2022).

### Cross-modal paired associative stimulation (cm-PAS) protocol

Cm-PAS had the same parameters as those described in Zazio et al. 2019. Participants were seated with their head on a chinrest to minimize head movements, at 57 cm distance from a computer monitor and wore noise-cancelling earphones playing white noise, to attenuate the click-sound of the TMS. They were asked to look at a red fixation cross superposed to a left-hand palm projected on the monitor in egocentric perspective. The hand was repeatedly paired with a TMS pulse over the right S1. The time interval between the visual-touch onset and the TMS pulse was 20 ms (cm-PAS_ISI-20_) or 100 ms (cm-PAS_ISI-100_), as in the original study (Zazio et al., 2019). TMS intensity was set at 150% of participants’ rMT, and 150 paired stimuli were delivered at a fixed frequency of 0.1 Hz. To ensure that participants were paying attention to visual-touch stimuli, they were asked to detect rare events during the cm-PAS: in 15 randomly-presented trials, the visual-touch frame was presented twice, and the TMS pulse was paired with the defined ISI (i.e., 20 or 100 ms) on the second touch (**Figure 3A**). Participants were asked to press a button on a computer keyboard with their right hand every time they detected the double visual-touch trials. The number of correct detections were recorded. The actual timing of visual-touch stimuli was checked using a photodiode. Trials randomization and timing of the stimuli were presented under computer control using the software E-Prime (2.0, Psychology Software Tool, Inc.).

**Figure 3.**
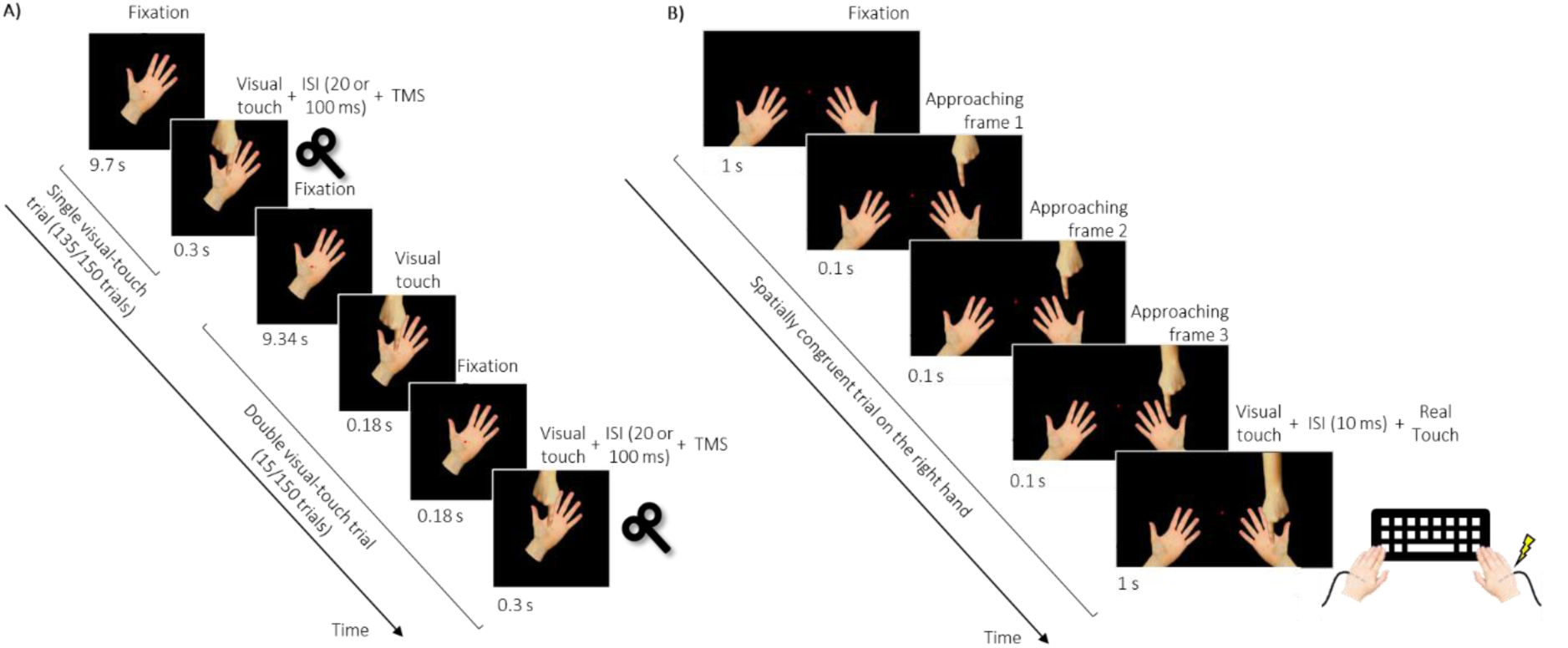
Schematic representation of the experimental paradigms. A) Cross-modal PAS (cm-PAS) trials: a single visual-touch trial followed by a double visual-touch trial. In single visual-touch trials (n=135), a fixation frame depicting a left-hand palm was presented for 9.7 s, followed by a visual-touch frame for 300 ms in which a hand in an allocentric perspective overlaps the hand palm in an egocentric perspective. After that, a new trial began without interruption, thus giving the illusion of an apparent motion of the hand touching the hand palm. In double visual-touch trials (rare events, n=15), the fixation frame was presented for 9.34 s, followed by a visual touch frame (180 ms), a fixation frame (180 ms) and a second visual touch frame (300 ms), thus giving rise to an apparent motion of a double touch on the hand palm. In both trial types, the TMS pulse was delivered after the visual touch trial lasting 300 ms, with a time interval of 20 ms (cm-PAS_ISI-20_) or 100 ms (cm-PAS_ISI-100_), in counterbalanced sessions. Participants were asked to maintain fixation on the red asterisk in the center of the palm in the same location where the visual touch occurred. B) Example of a spatially congruent trial in the visuo-tactile spatial congruity task. A fixation frame was presented for 1 s, depicting a left hand and a right hand in an egocentric perspective. Then, 3 approaching frames of 100 ms each were presented, showing the hands as in the fixation frame and another hand appearing from the upper part of the screen getting closer to the right hand. Finally, a visual touch stimulus was presented for 1 s, in which the hand in an allocentric perspective overlaps the hand palm in an egocentric perspective. The whole sequence within each trial gives the impression of an apparent motion terminating with a visual touch on the hand. Ten ms after the visual touch trial, a real touch was delivered on the participants’ right hand-palm. The same trial structure applied to spatially congruent trials on the left side and to spatially incongruent trials. In unimodal tactile-only trials, participants were shown with the two hands in an egocentric perspective, and only a tactile stimulus was delivered (thus, participants would have to provide a response, which was independent from visuo-tactile integration).

### Visuo-tactile spatial congruity (VTSC) task

The VTSC task was adapted from (Bolognini et al., 2014). Participants were seated in the same position as during the cm-PAS, and were asked to look at a red fixation cross at the center of a computer monitor while they were presented pictures of left- and right-hand palms in egocentric perspective. In visuo-tactile trials (n=120), another hand appeared from the upper part of the screen, touching either the left or the right-hand palm. Ten ms after the visual touch onset, a tactile stimulus was delivered either on the left or on the right-hand palm by means of small solenoid tappers. Therefore, the real and virtual hands were not in the same position, as in previous versions of the VTSC task (Bolognini, Miniussi, et al., 2013; Bolognini et al., 2014). The tactile stimulus was either spatially congruent (n=60) or incongruent (n=60) to the visual-touch stimulus, in random order. Participants were asked to report the location of the tactile stimulus, i.e., on the left or on the right hand, as fast and accurately as possible by pressing one of two buttons on a computer keyboard. Moreover, unimodal tactile-only (n=60) trials were presented as a check that participants were responding to tactile stimuli; finally, catch trials were provided (n=20) consisting of a brief change in the color of the fixation asterisk from red to green, to control for participants’ attention to the visual stimuli (see exclusion criteria paragraph and **Figure 2**). Further details on the timing of the visual stimuli are reported in **Figure 3B**. Before starting the experimental block, participants underwent a few practice trials, performed before the cm-PAS only. The VTSC total duration was approximately 8 min. The actual timing of visual-touch stimuli has been checked using a photodiode.

Both reaction RTs and accuracy (percentage of correct trials) were recorded. For each participant and each trial type, trials with RTs exceeding ± 2 SD were marked as outliers and discarded, after transforming the data in case of non-normality of the distribution (see Statistical Analysis for the normality assessment). For RTs, the difference between incongruent and congruent trials (ΔRTs_incong-cong_) was considered as the dependent variable.

Trials randomization and timing of the stimuli were presented under computer control using the software E-Prime (version 2.0, Psychology Software Tool, Inc.).

### Two-point discrimination task (2-PDT)

The 2-PDT was used to assess tactile acuity and was administered before and after cm-PAS. The procedure was adapted from (Zazio et al., 2019) and (L. K. Case et al., 2016; L. Case et al., 2017). Participants were blindfolded with their left arm relaxed on the desk, while an experimenter touched them on the thenar eminence of the left-hand palm with either one or two plastic tips, using an aesthesiometer, in a 2-alternative forced-choice task. In a range between 3 and 15 mm, 13 distances have been tested in descending blocks of 6 trials, each block comprising 3 trials with 1 tip and 3 trials with 2 tips, in random order. This procedure was repeated 2 times for a total of 12 trials for each distance tested (total number of trials: 156). A brief break was provided at the end of each repetition or if requested by the participants. A first experimenter, who was blind to the experimental session, was trained to deliver tactile stimuli with a frequency of approximately 0.5 Hz and to always apply the same pressure. Participants were asked to verbally respond (either ‘one’ or ‘two’) just after they had felt the touch, while a second experimenter recorded their responses on a computer. To ensure that participants had understood the task and to make them familiar with the sensations, before starting the experimental procedure, a few examples of touches with either one tip or two tips with a 15 mm distance were delivered, and feedback on their accuracy was provided by the experimenter. The 2-PDT estimated total duration was approximately 10 min.

Performance on the 2-PDT was evaluated according to the signal detection theory (Green & Swets, 1966), which disentangles the contribution of perceptual sensitivity (*d’*) and response bias (*c*). Specifically, we will consider as a dependent variable the sensory threshold, defined as the distance in mm corresponding to 50% of performance. The sensory threshold was estimated by fitting a logistic function to *d’* values (transformed to fit in a range between 0 and 1; (R Core Team, 2022). The fitting was done in R using the ‘fitting generalized linear models” (glm function, binomial family). If a participant showed negative values in the sensory threshold, she/he was discarded from this analysis.

## Preregistered Statistical Analysis

Statistical analysis was performed in Jamovi (v. 2.3.21; Study Design Template in **Table 1**). Statistical significance was set at p < 0.02.

I. First, BPD patients and HCs were compared in cognitive empathy levels obtained from the QCAE by means of a one-tailed Yuen’s trimmed mean t-test, a robust version of the t-test for independent samples (Mair & Wilcox, 2020).
II. Second, as a positive control, tactile acuity sensory threshold was analyzed in HCs using a repeated-measures analysis of variance (rm-ANOVA) with time (Pre_cm-PAS_, Post_cm-PAS_) and ISI (ISI-20, ISI-100).
III. Moreover, to assess the effects of cm-PAS on VTSC task performance in HCs, a rm-ANOVA with factors Time (Pre_cm-PAS_, Post_cm-PAS_) and ISI (ISI-20, ISI-100) was performed on ΔRTs_incong-cong_.
IV. Finally, the two groups were compared in the effects induced by the cm-PAS on tactile acuity sensory threshold and VTSC task performance (RTs). The analyses consisted of a 2×2×2 mixed-design ANOVA with between factor Group (BPD, HCs) and within factors Time (Pre_cm-PAS_, Post_cm-PAS_) and ISI (ISI-20, ISI-100).

Normality of continuous data was assessed by means of Shapiro-Wilk test. Whenever deviating from normality, data were transformed according to the commonly used transformations for continuous data, as there is no robust version of rm- or mixed-design ANOVA: square root, i.e., √(raw data); base-ten logarithmic, i.e., log10(raw data), and (c) inverse transformation, i.e., 1/(raw data). To account for possible negative values, as well as values between 0 and 1, when applying these transformations, we added a constant to the raw data values, thus anchoring the minimum of our distribution(s) to 1 (Osborne & Carolina, 2016). Then, the transformation showing the best fit to a normal distribution was assessed as the shortest Euclidean distance from a normal distribution, according to Cullen-Frey graphs and the ‘fitdistrplus’ package in R (Delignette-Muller & Dutang, 2015); https://cran.r-project.org/web/packages/fitdistrplus/index.html). If none of these transformations made the data distribution close enough to normality, i.e., the transformed distribution presented values of an excess kurtosis (beyond 3 ± 2) and square of skewness > 1 on Culley Frey graph (George & Mallery, 2019), we proceeded with non-parametric tests.

While the analyses described above were the ones that drove the interpretation of results and the conclusions, results were also supported by equivalent tests using Bayesian statistics, performed in JASP (v. 0.17.1.0).

## Results

### Clinical assessment

The mean scores of the clinical assessments of individuals with BPD in the final sample are shown in Table 2. Overall, individuals with BPD exhibited moderate BPD symptoms. Patients also showed elevated scores, indicating marked symptom severity, in depression, impulsivity, alexithymia, global symptom severity, and trauma history. More than 90% of participants reported engaging in self-harm behaviors. Regarding substance use, 20% of individuals with BPD reported current or past alcohol abuse, and 30% reported current and past drug abuse. The mean clinical scores of the full sample of patients enrolled in this study are presented in Table S1. No substantial differences in symptom severity were observed between the final and full enrolled samples.

**Table 2.** Test scores at the clinical assessment of patients with borderline personality disorder (BPD), considering the final sample of 24 patients. N indicates the number of participants who completed the questionnaires.

|  | N | Mean | ± | SE |
| --- | --- | --- | --- | --- |
| Zanarini rating scale for Borderline Personality Disorder (ZAN-BPD) | 21 | 9.62 | ± | 0.84 |
| Baratt Impulsiveness Scale (BIS-11) | 21 | 74.24 | ± | 1.99 |
| Beck Depression Inventory (BDI_II) | 21 | 34.33 | ± | 2.65 |
| Toronto Alexithymia Scale (TAS) | 20 | 64.25 | ± | 3.01 |
| Symptom Checklist-90-R (SCL90) | 21 | 2.14 | ± | 0.17 |
| Inventory of Interpersonal Problems (IIP) | 21 | 2.22 | ± | 0.14 |
| Childhood Trauma Questionnaire (CTQ) | 20 | 55.6 | ± | 4.51 |
| Attachment Style Questionnaire (ASQ) | 19 |  |  |  |
| *Confidence* |  | 21.84 | ± | 1.33 |
| *Discomfort with Closeness* |  | 41 | ± | 16.11 |
| *Relationships as secondary* |  | 16.11 | ± | 1.49 |
| *Need for Approval* |  | 33.63 | ± | 1.65 |
| *Preoccupation with Relationships* |  | 36.26 | ± | 2.11 |
| Inventory of statements about self-injury (ISAS) | 20 |  |  |  |
| *Presence of self-injury (%)* |  | 95% |  |  |

### Preregistered hypotheses

Bayesian analyses were interpreted according to the classification scheme for reporting results from JASP (Kelter, 2020).

#### I. Cognitive empathy levels: HCs vs. individuals with BPD

The comparison between patients with BPD and HCs on cognitive empathy scores from the QCAE showed a trend toward significance in the expected direction when applying the preregistered significance threshold of p = 0.02 (t = 1.95, p = 0.031; **Figure 4**). While the preregistered frequentist analysis employed a robust Yuen’s trimmed mean t-test, complementary Bayesian analyses were performed using a standard independent samples t-test in JASP, as no robust Bayesian alternative is currently available. Complementary Bayesian analysis yielded a Bayes Factor (BF_10_) of 2.82, providing anecdotal evidence in favor of the preregistered hypothesis. The average scores of the final sample are presented in Table 3.

**Figure 4.**
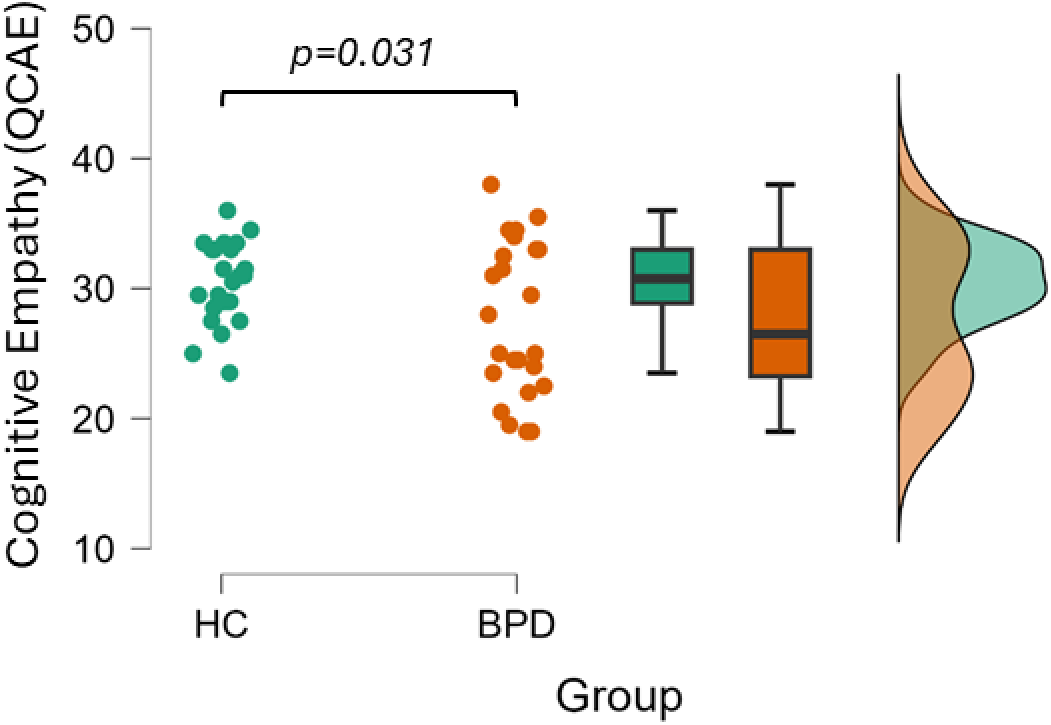
Raincloud plot showing raw data (dots), boxplots, and kernel density distributions for the final sample of healthy controls (HC, green, N = 24) and patients with borderline personality disorder (BPD, orange, N = 24) on the cognitive empathy scores of the Questionnaire of Cognitive and Affective Empathy (QCAE). The p-value corresponds to the one-tailed Yuen’s trimmed mean t-test comparing HCs and patients with BPD (hypothesis I).

**Table 3.**
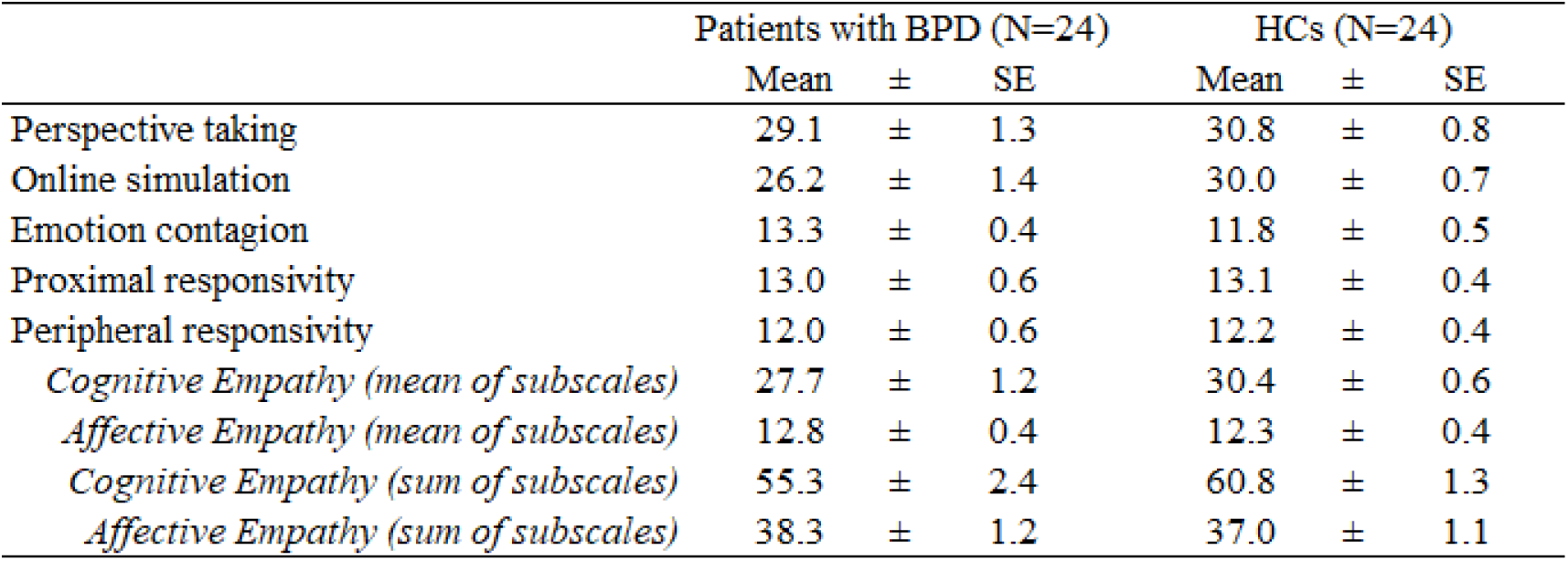
Mean and standard error (SE) for scores obtained at the Questionnaire for Cognitive and Affective Empathy in the final sample of patients with borderline personality disorder (BPD) and healthy controls (HCs).

| | Patients with BPD ($N=24$) | | | HCs ($N=24$) | | |
| --- | --- | --- | --- | --- | --- | --- |
| | Mean | ± | SE | Mean | ± | SE |
| Perspective taking | 29.1 | ± | 1.3 | 30.8 | ± | 0.8 |
| Online simulation | 26.2 | ± | 1.4 | 30.0 | ± | 0.7 |
| Emotion contagion | 13.3 | ± | 0.4 | 11.8 | ± | 0.5 |
| Proximal responsivity | 13.0 | ± | 0.6 | 13.1 | ± | 0.4 |
| Peripheral responsivity | 12.0 | ± | 0.6 | 12.2 | ± | 0.4 |
| *Cognitive Empathy (mean of subscales)* | 27.7 | ± | 1.2 | 30.4 | ± | 0.6 |
| *Affective Empathy (mean of subscales)* | 12.8 | ± | 0.4 | 12.3 | ± | 0.4 |
| *Cognitive Empathy (sum of subscales)* | 55.3 | ± | 2.4 | 60.8 | ± | 1.3 |
| *Affective Empathy (sum of subscales)* | 38.3 | ± | 1.2 | 37.0 | ± | 1.1 |

We further conducted an exploratory analysis on the full sample of participants tested (35 HCs and 36 patients with BPD), as the preregistered analysis involved a smaller subset because of preregistered exclusion criteria applied to other measures. In this enlarged sample, the group difference reached significance (t = 2.67, p = 0.011), suggesting that the smaller sample size in the preregistered analysis had reduced statistical power (**Figure S1**). Complementary Bayesian analysis yielded a BF_10_ of 13.9, providing strong evidence that supports the expected group differences. Table S2 presents the average scores of the full sample.

To facilitate a comparison with the previous study by our group (Zazio et al., 2024), Table 3 and Table S2 also report the mean and standard error calculated on the sum of the “perspective taking” and “online simulation” subscales, in addition to the mean values considered in the preregistered analyses.

#### II. Plasticity effects of cm-PAS on tactile acuity in HCs (positive control)

The sensory threshold of tactile acuity was not normally distributed (W = 0.76, p < 0.001), and neither of the planned transformations brought the data distribution within the preregistered ranges for skewness and kurtosis. Therefore, as preregistered, we proceeded with non-parametric analyses, specifically using the Aligned Rank Transform (ART) procedure, which accounts for the repeated-measures structure. Results from the ART ANOVA showed no significant main effects of time (F(1,84) = 0.23, p = 0.633) or ISI (F(1,84) = 0.12, p = 0.731), or a significant time × ISI interaction (F(1,84) = 0.01, p = 0.921). While the preregistered analysis was performed using a robust ART ANOVA to account for non-normality and repeated measures, complementary Bayesian analyses were conducted using a standard Bayesian rm-ANOVA in JASP, as no direct Bayesian analog of the ART procedure is currently available. Consistent with the non-significant frequentist results, the Bayesian analysis provided strong evidence in favor of the null hypothesis for the time × ISI interaction (BF_10_ = 0.06) and anecdotal evidence for the null hypothesis for the main effects (ISI: BF_10_ = 0.61; time: BF_10_ = 0.31; **Figure 5**).

**Figure 5.**
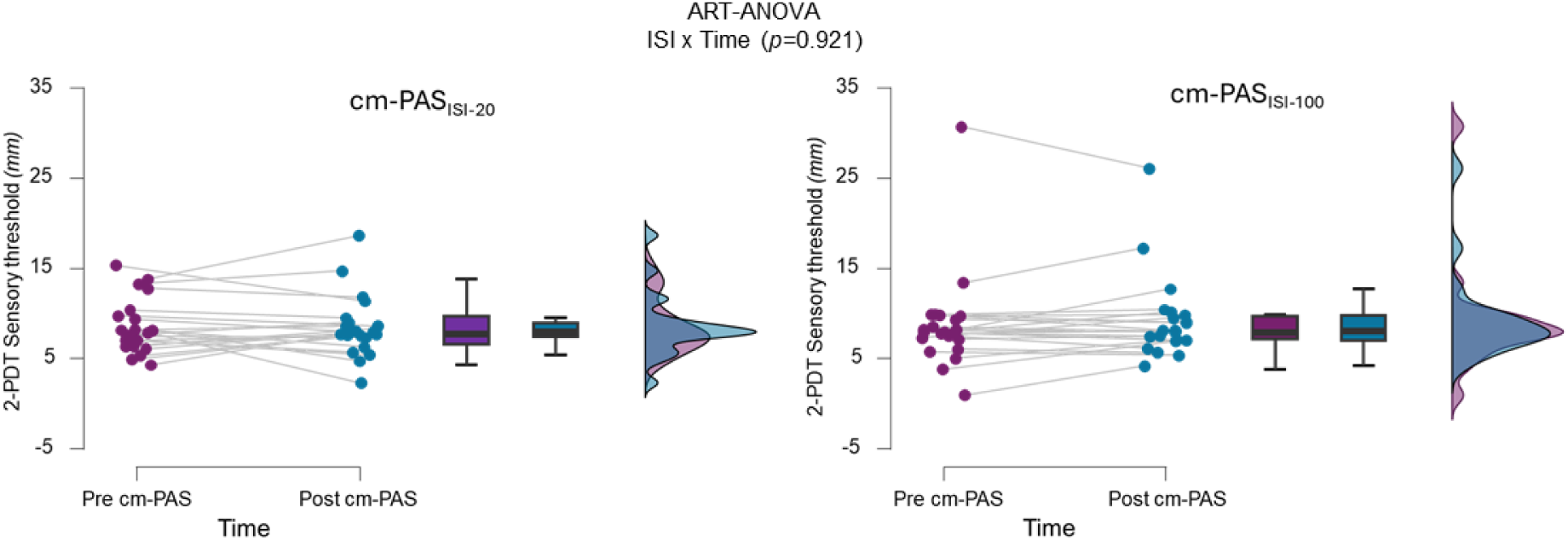
Raincloud plot showing raw data (dots), boxplots, and kernel density distributions for the final sample of healthy controls on tactile acuity in the two-point discrimination task (2-PDT), expressed as sensory threshold in mm derived from d′ values, before and after the cross-modal paired associative stimulation (cm-PAS) protocol. The preregistered repeated-measures analysis revealed no significant time × inter-stimulus interval (ISI) interaction (hypothesis II), nor main effects of time (pre, post) or ISI (ISI-20, ISI-100).

**Figure 6.**
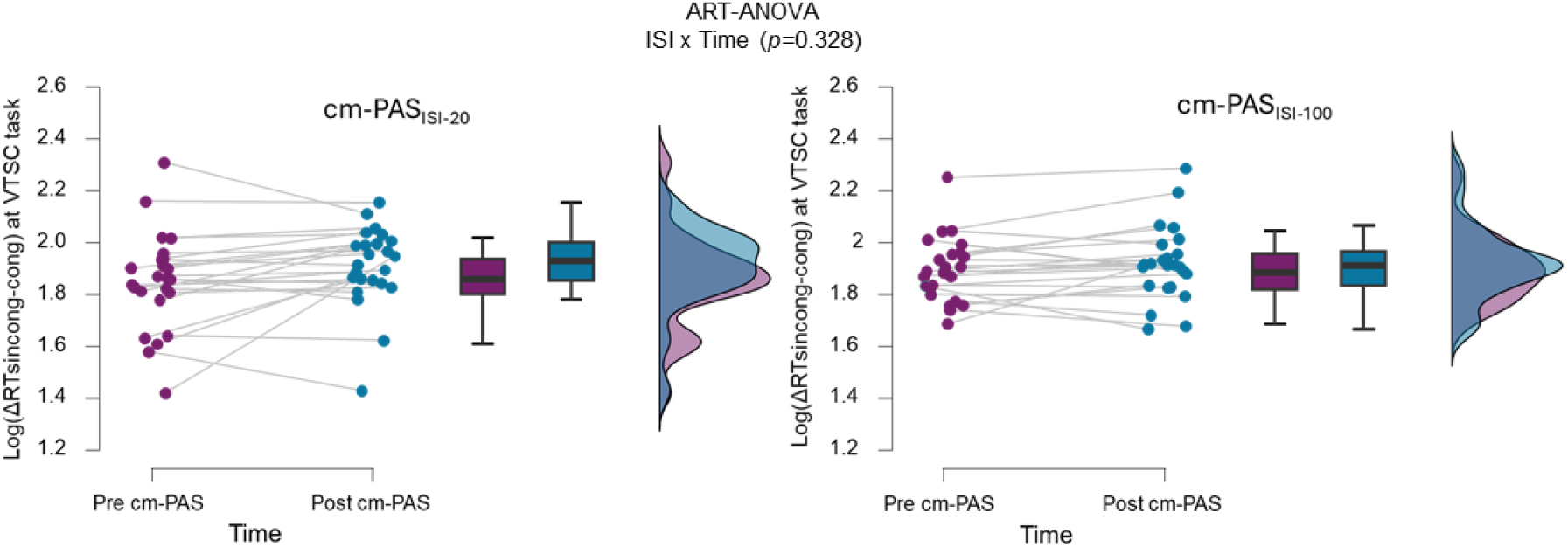
Raincloud plot showing raw data (dots), boxplots, and kernel density distributions for the final sample of healthy controls on the interference effect in the visuo-tactile spatial congruency (VTSC) task, before and after the cross-modal paired associative stimulation (cm-PAS) protocol. The interference effect is expressed as the difference between reaction times in incongruent vs. congruent trials (log-transformed ΔRTsincong-cong). The preregistered repeated-measures analysis revealed no significant time × inter-stimulus interval (ISI) interaction (hypothesis III), nor a main effect of ISI (ISI-20, ISI-100), but a significant main effect of time (pre, post), indicating overall longer RTs after cm-PAS.

**Figure 8.**
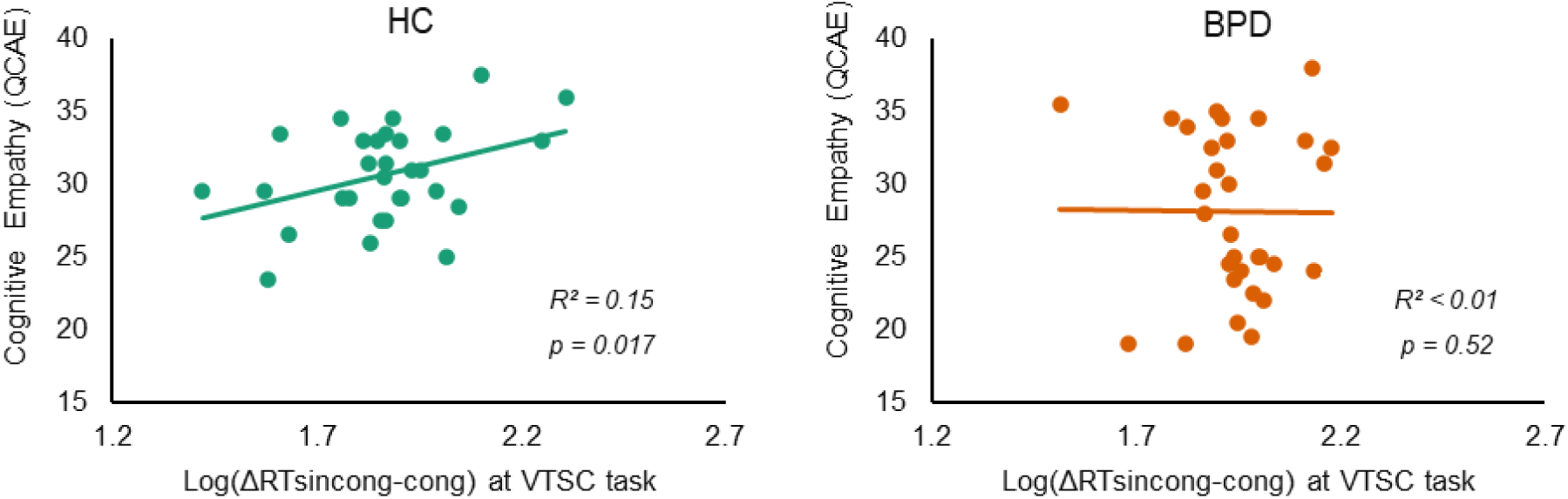
Exploratory analysis of the relationship between cognitive empathy scores on the Questionnaire for Cognitive and Affective Empathy (QCAE) and the interference effect at baseline in the visuo-tactile spatial congruency (VTSC) task. The interference effect is expressed as the difference between reaction times in incongruent vs. congruent trials (log-transformed ΔRTsincong-cong). Data are shown for healthy controls (HC, green, left panel) and patients with borderline personality disorder (BPD, orange, right panel). P-values indicate within-group correlations, examined in the presence of a significant empathy × group interaction.

A complementary, non-preregistered analysis was conducted to further characterize the interindividual variability in cm-PAS responsiveness. Given that cm-PAS ISI-20 is expected to lower the sensory threshold, enhancing tactile acuity (Maddaluno et al., 2020; Zazio et al., 2019), the participants who exhibited modulation on the 2-PDT sensory threshold 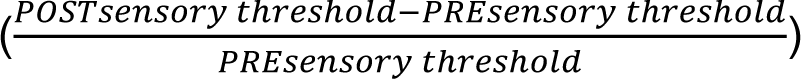 smaller than −0.05 (i.e., −5%) were considered as “PAS responders.” This cutoff was chosen to avoid considering participants presenting negligible modulation, likely owing to measurement variability (for a similar procedure, see Arrigoni et al., 2025). According to this criterion, 10 of 22 individuals (45%) were classified as responders.

#### III. Plasticity effects of cm-PAS on TaMS functioning in HCs

The ΔRTs_incong-cong_ at the VTSC task was not normally distributed (W = 0.89, p < 0.001); among the planned transformations, the base-ten logarithm brought the distribution within the preregistered range of skewness and kurtosis. The rm-ANOVA revealed a significant main effect of time (F(1,23) = 6.46, p *=* 0.018), indicating that overall, ΔRTs_incong-cong_ was longer after the cm-PAS. However, neither a significant main effect of ISI (F(1,23), p = 0.158) nor a significant time × ISI interaction (F(1,23) = 1.0, p = 0.328) was observed. The Bayesian approach yielded a BF_10_ of 0.3 for the time × ISI interaction, indicating moderate evidence for the null hypothesis, consistent with the non-significant outcome observed in the frequentist analysis. Regarding the main effect of time, the Bayesian analysis provided no evidence favoring either the null or alternative hypothesis (BF_10_ = 1), whereas the main effect of ISI offered moderate/anecdotal evidence in favor of the null hypothesis (BF_10_ = 0.33).

#### IV. Plasticity effects of cm-PAS: HCs vs. individuals with BPD

Given that the cm-PAS ISI-20 was ineffective in inducing plastic changes in HCs in either the sensory threshold measured with 2-PDT or VTSC performance, a comparison between HCs and individuals with BPD on plasticity effects (hypothesis IV) was not performed. Nonetheless, an exploratory responder classification revealed an identical proportion of responders in the BPD group compared with HCs (45%), based on the same ≥ 5% modulation criterion on tactile acuity reported previously.

## Exploratory analyses

Exploratory analyses were conducted to provide complementary insights beyond the preregistered analyses. Specifically, these analyses aimed to further characterize the effects of cm-PAS in HCs, compare baseline measures in HCs and patients with BPD, and examine their potential associations with individual differences in cognitive empathy. For each exploratory analysis, the sample was defined so as to maximize statistical power. Accordingly, when the effects of cm-PAS were not under investigation, participants who did not complete the second experimental session were also included. These analyses were not preregistered and should be interpreted with caution; however, they offer valuable insights that may inform hypotheses for future studies. A conventional significance threshold of p = 0.05 was applied.

### Characterization of the effects of cm-PAS in HCs

Within the exploratory framework, a series of analyses was conducted to further characterize the effects of cm-PAS by examining potential modulatory factors and sources of interindividual variability in HCs. Specifically, we explored (i) the possible attenuation of plasticity effects over time by accounting for task order in the assessment of the effects of cm-PAS and (ii) the impact of the number of trials used to assess tactile acuity on the stability of threshold estimates.

#### (i) Effects of cm-PAS on tactile acuity accounting for task order

In the present study, the participants completed two behavioral tasks after cm-PAS, unlike the previously used protocol, which only assessed tactile acuity (Zazio et al., 2019). The order of task administration was counterbalanced: half of the participants performed the 2-PDT first, followed by the VTSC, while the remaining participants completed the tasks in the opposite order. Because both tasks were administered within approximately 25 min after the end of cm-PAS (i.e., the expected window for plasticity effects), this procedural difference was not expected to substantially affect the outcomes. Nonetheless, to rule out the possibility of attenuated plasticity over time, we conducted an exploratory analysis accounting for the order effect by running an ART ANOVA with within-subject factors time (pre, post) and ISI (ISI-20, ISI-100) and between-subject factor order (VTSC-first, 2-PDT-first). If the PAS-induced plasticity effects on tactile acuity rapidly dissipated over time, we would expect the order to significantly interact with ISI and time; however, this was not the case (F(1,80) = 0.07, p = 0.787). A complete report of these results is provided in Table S3. This result is consistent with the absence of the effect of cm-PAS at the group level.

#### (ii) Potential impact of trial number in tactile acuity assessment

Another difference compared with the previous study from our group, in which the cm-PAS protocol was adapted, is the substantially lower number of trials used to assess tactile acuity in the 2-PDT (156 vs. 390 trials (Zazio et al., 2019)). This procedural difference raised the possibility that the reduced number of trials might have increased interindividual variability in sensory threshold estimates, potentially leading to a less reliable assessment of tactile acuity before and after cm-PAS. To address this issue, we reanalyzed data from the original study (Zazio et al., 2019) by estimating the sensory threshold using only the first repetition of the task, yielding a number of trials comparable to those employed in the present study (13 distances × 10 trials; 130 trials). We then assessed the agreement between sensory thresholds obtained from the full dataset (390 trials) and from the reduced subset (130 trials) using the intraclass correlation coefficient (ICC), a measure of reliability that captures agreement between repeated measurements and interindividual variability (Mcgraw, 1996) and that has been previously applied to evaluate the reliability of estimates obtained with different numbers of trials (Biabani, Farrell, Zoghi, Egan, & Jaberzadeh, 2018). The ICC was computed based on the baseline 2-PDT sensory thresholds from the first experiment (n = 16), using a two-way mixed-effects model, single measurement, and absolute agreement definition (ICC(3,1)). The analysis yielded an ICC of 0.81, indicating good reliability according to established guidelines (Koo & Li, 2016), thus suggesting that reducing the number of trials did not substantially compromise the reliability of the 2-PDT sensory threshold in this paradigm.

### Baseline comparison of tactile acuity and TaMS functioning: HCs vs. individuals with BPD

An exploratory analysis was conducted to investigate potential differences in the baseline tactile acuity and VTSC task between groups, which may inform future studies on the somatosensory domain in BPD. Focusing on baseline measures, only data from the first session of the 2-PDT and VTSC tasks were analyzed, and participants who did not complete the second session were also included to maximize statistical power.

For the 2-PDT sensory threshold, 27 HCs and 28 individuals with BPD were included. As the data were non-normally distributed (W = 0.42, p < 0.001) and could not be normalized via transformation, a two-tailed Mann-Whitney test was applied, revealing no significant difference between groups (U = 360, p = 0.471).

For the VTSC task, 30 HCs and 31 individuals with BPD were included. A logarithmic transformation of ΔRTsincong-cong achieved an acceptable distribution, allowing a two-tailed Student’s t-test, which did not show a significant group difference (t = -1.78, p = 0.081). Accuracy differences between incongruent and congruent trials (ΔRTsincong-cong) were also explored using a non-parametric test due to non-normality, with no significant differences observed (U = 355, p = 0.102). Moreover, we also explored VTSC performance in terms of accuracy, and specifically examining on the difference in accuracy between incongruent and congruent and incongruent trials (i.e., ΔACCincong-cong), applying a non-parametric test due to the non-normal distribution of the data. No significant differences emerged in the ΔACCincong-cong (U=355, p=0.102).

Overall, no significant differences emerged between HCs and BPD participants in either tactile acuity or TaMS functioning.

### Relationship between cognitive empathy levels and VTSC performance

The effect of VTSC interference indexes the extent to which observed touches influence tactile processing and has been proposed to capture the engagement of somatosensory mirroring mechanisms. Building on this framework, we examined the relationship between individual differences in empathic abilities and VTSC performance as described previously (Bolognini, Miniussi, et al., 2013; Bolognini et al., 2014).

As in the previous exploratory analysis, only baseline VTSC measures (i.e., ΔRTs_incong-cong_) from the first session were analyzed here, including also participants who did not complete the second session. We ran an ANCOVA with the log-transformed ΔRTs_incong-cong_ as the dependent variable, group (HCs, BPD) as a between-subject factor, and the cognitive empathy score from the QCAE as a covariate.

ANCOVA revealed a significant main effect of group (F(1,57) = 5.99, p = 0.017), a significant main effect of cognitive empathy (F(1,57) = 4.59, p = 0.037), and a significant group × cognitive empathy interaction (F(1,57) = 4.77, p = 0.033). Given the presence of an interaction, we focused on this effect by examining the simple correlations within each group. Consistent with the directional hypothesis that higher cognitive empathy would be associated with stronger visuo-tactile interference (i.e., larger ΔRTs_incong-cong_), a significant positive correlation was observed in HCs (r = 0.39, p = 0.017). A complementary Bayesian analysis yielded a BF_10_ of 3.8, indicating moderate evidence for the predicted association. In contrast, no significant relationship was observed in individuals with BPD (r = −0.01, p = 0.520) with a BF_10_ of 0.2, providing moderate evidence of no positive association.

## Discussion

The present study investigated the empathic abilities and integrity of plasticity mechanisms underlying visuo-tactile mirroring phenomena in patients with BPD, employing a previously used TMS protocol that acts on Hebbian associative plasticity and exploits the mirror properties of the somatosensory system (i.e., cm-PAS; Zazio et al., 2019). The preregistered analyses did not fully support our hypotheses: the difference in cognitive empathy between patients with BPD and HCs did not reach the preregistered significance threshold, and the expected plasticity effects of cm-PAS in HCs were not replicated, precluding a direct comparison of plasticity mechanisms between groups. Despite these null confirmatory findings, the study provides a rigorous preregistered assessment of cognitive empathy and behavioral measures of tactile acuity and performance in a cross-modal task involving the TaMS. Moreover, exploratory analyses enabled the evaluation of the impact of potential modulatory factors on cm-PAS efficacy as well as the relationship between empathic dimensions and visuo-tactile processing.

The preregistered analyses focused on cognitive empathy levels and plasticity effects of cm-PAS. Regarding empathic abilities, previous studies have reported alterations in empathic abilities in patients with BPD, particularly in understanding others’ perspectives (i.e., cognitive empathy) rather than feeling others’ emotions (i.e., affective empathy) (Grzegorzewski et al., 2019; Zazio et al., 2024) (Harari et al., 2010; E. Martin, Blais, Albaret, Pariente, & Tallet, 2017). In the present sample of 24 participants per group, the comparison between patients with BPD and HCs in terms of cognitive empathy scores showed a trend in the expected direction (p = 0.031). Still, it did not meet the preregistered significance threshold (p < 0.02). Therefore, this effect could not be considered confirmatory.

Several converging observations suggest that this null confirmatory effect is more likely due to limited statistical power rather than to a substantive inconsistency with the preregistered hypothesis. First, the clinical profile of the present sample closely matched that of our previous study on patients with BPD in terms of symptom severity (Zazio et al., 2024), reflecting a similar recruitment context. Second, complementary Bayesian statistics favored the preregistered hypothesis, albeit with anecdotal evidence. Third, when considering an enlarged sample of 36 patients with BPD who met all diagnostic inclusion criteria but failed other exclusion criteria unrelated to empathy measures, the group difference reached statistical significance and was accompanied by strong Bayesian support for reduced cognitive empathy in patients with BPD compared with that in HCs. Finally, the mean cognitive empathy scores observed here closely align with those reported in our previous study, which revealed a significant group difference (Zazio et al., 2024). Collectively, these results suggest that cognitive empathy alterations are relevant in BPD but are not uniformly expressed across individuals with this diagnosis. Last but not least, self-report measures of empathy may be limited in patients with BPD due to variability in introspective insight, emotional dysregulation, and context-dependent response styles. It should be noted that patients with BPD can usually empathize adeptly, but in interpersonal/social situations in which emotional arousal is higher and attention span is limited the ability is more impaired (Bateman & Fonagy, 2004),This heterogeneity can reduce effect sizes, highlighting the need for complementary behavioral measures and larger, adequately powered samples to reliably detect group-level differences. Nevertheless, despite some limitations, self-report instruments are widely used and accepted in the literature due to their practicality and their ability to assess the subjective experience of empathy.

The cm-PAS manipulation in HCs, designed as a positive control, did not induce significant changes in tactile acuity, as assessed using 2-PDT, indicating a failure to replicate previously reported plasticity effects in the somatosensory system (Maddaluno et al., 2020; Zazio et al., 2019). Moreover, the VTSC task performance did not provide evidence of cm-PAS-induced modulation of TaMS function. As a result, the effectiveness of cm-PAS could not be confirmed within the preregistered framework, precluding the planned comparison of plasticity mechanisms between HCs and patients with BPD.

To better contextualize these null findings, we explored several potential sources of variability. First, we tested whether the sequential administration of the two tasks after the cm-PAS (i.e., the 2-PDT and VTSC in a counterbalanced order) might have attenuated the expected effect when the 2-PDT was performed. This exploratory analysis provided no evidence of an order effect, indicating that the lack of cm-PAS modulation did not depend on whether 2-PDT was administered immediately after stimulation or approximately 10 min after its completion.

Several methodological considerations warrant further investigation. Although the cm-PAS protocol itself was identical to that used in an earlier study (the same monophasic stimulator, stimulation parameters, and timing), other aspects differed from the original design. First, 2-PDT was administered in a substantially reduced number of trials (156 vs. 390), which may have increased the measurement variability. A reanalysis of previously published data (Zazio et al., 2019) using a subset of trials showed good agreement between tactile acuity thresholds derived from the reduced and full datasets, as indexed by the ICC, suggesting that the lower trial number is unlikely to have substantially compromised the reliability of tactile acuity in the present study and, therefore, to have accounted for the absence of cm-PAS effects. Additionally, the stimulation site in S1 was localized according to more recent literature (Holmes & Tamè, 2019; Holmes et al., 2019), placing the target 2 cm lateral and 0.5 cm posterior to the primary motor cortex hotspot, rather than 2 cm posterior as in previous studies (Gorgoni, Ferlazzo, Atri, Lauri, & Ferrara, 2015; Litvak et al., 2007; A. Wolters et al., 2005). This updated targeting is supported not only by systematic reviews and meta-analytic evidence but also by experimental TMS studies specifically targeting S1, providing converging evidence for this localization. Although individual magnetic resonance images were not available in the present study, this approach was derived from the functionally identified primary motor cortex hotspot, allowing for the adjustment of the stimulation site based on each participant’s functional mapping. Even if this method does not precisely correspond to S1 in every individual, as it reflects the average across studies, the relatively high stimulation intensity used during cm-PAS (i.e., 150% rMT) likely generated a sufficiently broad electric field to encompass S1 across participants.

In this study, of 22 healthy individuals, only 10 (45%) could be classified as “PAS responders” on tactile acuity, with an identical proportion observed in patients with BPD. However, this proportion should be interpreted with caution, as a similar distribution would be expected if post-pre differences were randomly distributed around zero. Accordingly, the present data do not provide strong support for a categorical distinction between responders and non-responders, but rather highlight substantial inter-individual variability in cm-PAS effects. Such high variability in plasticity-inducing TMS protocols is well documented in the literature, extending beyond the PAS paradigms (Huang et al., 2017; Magnuson et al., 2023). Several studies have proposed that individuals may differ substantially in their propensity to exhibit measurable plasticity effects, leading to a distinction between “responders” and “non-responders” (Nettekoven et al., 2015). This variability has prompted a growing interest in approaches that tailor stimulation to individual neurophysiological characteristics to enhance the reliability of induced plasticity (e.g., accounting for functional connectivity patterns (Bagattini et al., 2021) or timing TMS pulses to ongoing neural oscillations (Zrenner & Ziemann, 2024)). Within this broader context, the absence of a cm-PAS effect in the present sample may reflect the well-known variability in plasticity-inducing TMS protocols, rather than provide definitive evidence against the effectiveness of cm-PAS.

At baseline, the results revealed no significant differences in tactile acuity or VTSC interference between the HCs and participants with BPD. These findings align with those of previous studies that reported selective impairments in certain aspects of somatosensation, such as nociception and tactile sensitivity, while sparing others, including tactile acuity and TaMS functioning, when assessed using VTSC performance as a behavioral marker of cross-modal deficits at the level of TaMS (Cruciani et al., 2023; Pavony & Lenzenweger, 2014; Zazio et al., 2024). Among HCs, exploratory analyses revealed that individuals with higher cognitive empathy experienced greater interference in the VTSC task, consistent with prior work suggesting that stronger perspective-taking ability amplifies interference when observing others being touched while reporting the location of one’s own tactile stimulation (Bolognini, Miniussi, et al., 2013; Bolognini et al., 2014). Notably, the present study provides the first evidence that such a relationship can be detected at baseline without prior experimental modulation of S1 activity. For transparency, a previous study from our group, which employed the same task, found no significant correlation between cognitive empathy and VTSC performance, a result that may be because of the small sample size (approximately 30% smaller) (Zazio et al., 2024).

In contrast, this association was absent in BPD, as indicated by the significant interaction between the group and empathy variables. Such an association would still be expected if the VTSC performance captured the ability to resonate with others’ touch, despite the lower average cognitive empathy typically observed in this group. The lack of an association cannot be explained by restricted ranges or floor effects. One possibility relies on the limitations of a self-report measure of empathy in individuals with BPD discussed above. Alternatively,, the results may suggest a more complex relationship, in which TaMS functioning, as measured by VTSC performance, does not directly map onto cognitive empathy. This pattern aligns with previous findings from the visuomotor mirroring literature, which highlights the complex interplay between individuals’ empathic abilities and the magnitude of neurophysiological indices assessing mirror neuron system recruitment according to the features of the observed action (Bekkali et al., 2021; Guidali et al., 2025; Guidali, Picardi, Franca, Caronni, & Bolognini, 2023; Zhao, Li, Xiang, & Liu, 2024). Similarly, within the TaMS, higher-order cognitive processes involved in the elaboration of observed touch, such as the integration of social and tactile cues, can modulate visuo-tactile interference. This modulation may contribute to non-significant or atypical correlational patterns in a clinical population with impaired empathy, such as patients with BPD, highlighting the need for further studies to understand these dynamics.

## Conclusions

The present study provides a rigorous evaluation of cognitive empathy, tactile processing, and putative plasticity mechanisms within the TaMS in patients with BPD. Although the preregistered hypotheses were not supported, owing to the lack of confirmatory group differences in cognitive empathy and the absence of plasticity effects induced by cm-PAS, the study offers important methodological and empirical contributions. The null plasticity findings limited the planned group comparison but also highlighted the inherent variability in inducing S1 plasticity across individuals.

Overall, tactile acuity and visuo-tactile interference appear to be preserved in patients with BPD; however, the association between cognitive empathy and TaMS functioning observed in healthy individuals is disrupted in patients. The absence of this relationship in BPD suggests that the link between empathy and tactile mirroring depends on additional processes that shape visuo-tactile integration. Collectively, these findings emphasize the importance of adequately powered samples, reliable stimulation protocols, and multimodal approaches for clarifying the role of the TaMS in BPD. Future studies are needed to determine whether and how empathic abilities interact with somatosensory processing and to establish the conditions under which plasticity in the S1 can be reliably modulated.

## Authors’ contribution

AZ: conceptualization, investigation, data curation, methodology, formal analysis, visualization, writing -original draft, project administration, funding acquisition; supervision; GG: conceptualization, formal analysis, writing -original draft; CML: investigation, formal analysis, writing -review and editing; CM: investigation, writing -review and editing; ED: investigation, writing -review and editing; SM: investigation, writing -review and editing; BB: writing -review and editing; RR: resources, writing -review and editing; NB: conceptualization, writing -review and editing; MB: conceptualization, writing -original draft.

## Acknowledgements

We would like to thank Claudia Fracassi and Andrea Tranchina for their support in data collection.

## Data availability statement

The codes and the stimuli of the study are publicly available at https://osf.io/5hvpf/overview, while the data can be found at 10.5281/zenodo.18327056.

## Abbreviations’ list

2-PDT: Two-Point Discrimination Task
APB: Abductor Pollicis Brevis
ASQ: Attachment Style Questionnaire
BDI-II: Beck Depression Inventory II
BIS: Barratt Impulsiveness Scale
BPD: Borderline Personality Disorder
cm-PAS: Cross-Modal Paired Associative Stimulation
CTQ: Childhood Trauma Questionnaire
DSM-5: Diagnostic and Statistical Manual of Mental Disorders, Fifth Edition
HC: Healthy Control
ICC: Intraclass Correlation Coefficient
IIP: Inventory of Interpersonal Problems
ISAS: Inventory of Statements About Self-injury
ISI: Inter-Stimulus Interval
LTD: Long-Term Depression
LTP: Long-Term Potentiation
NMDA: N-methyl-D-aspartate
PAS: Paired Associative Stimulation
QCAE: Questionnaire of Cognitive and Affective Empathy
rMT: Resting Motor Threshold
RT: Reaction Times
S1: Primary Somatosensory Cortex
SCID-CV: Structural Clinical Interview for DSM Disorder -Clinician Version
SCID-PD: Structural Clinical Interview for DSM Personality Disorder
SCL-90-R: Symptoms Check-list 90 Revised
TaMS: Tactile Mirror System
TAS-20: Toronto Alexithymia Scale
TMS: Transcranial Magnetic Stimulation
TMS_SensQ: Self-report Questionnaire on Sensations induced by TMS
VTSC: Visuo-Tactile Spatial Congruity
ZAN-BPD: Zanarini rating scale for BPD

