## Supplementary Materials for "Cortical plasticity of the tactile mirror system in borderline personality disorder"

**Figure 1.** Raincloud plot showing raw data (dots), boxplots, and kernel density distributions for the final sample of healthy controls (HC, green, N=35) and patients with borderline personality disorder (BPD, orange, N=36) on the Cognitive Empathy scores of the Questionnaire of Cognitive and Affective Empathy (QCAE). The p-value corresponds to the one-tailed Yuen’s trimmed mean t-test.

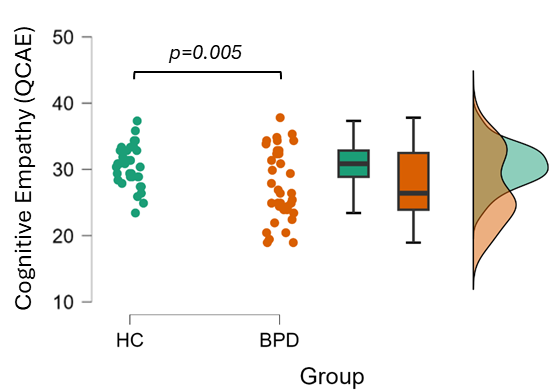

**Table S1.** Test scores at the clinical assessment of patients with borderline personality disorder (BPD), considering the full sample of 36 patients who were enrolled in the study. N indicates the number of participants who completed the questionnaires.

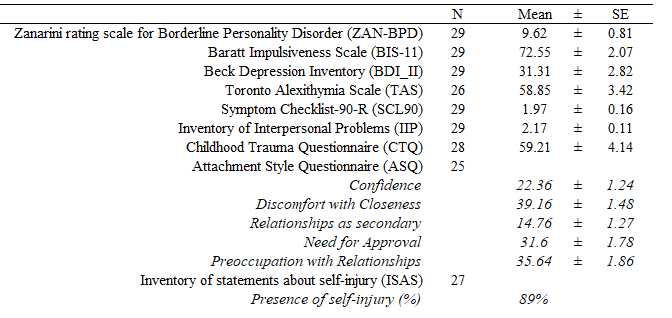

**Table S2**. Mean and standard error (SE) for scores obtained at the Questionnaire for Cognitive and Affective Empathy (QCAE) in the full sample of patients with Borderline Personality Disorder (BPD) and healthy controls (HCs).

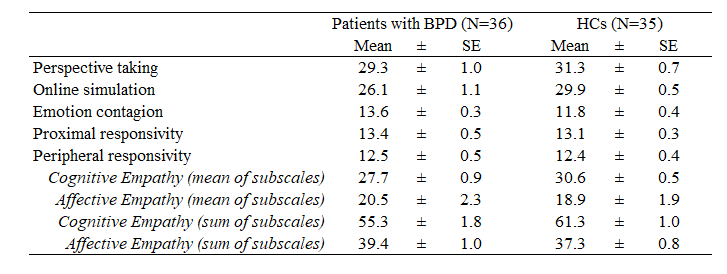

**Table S3**. Cm-PAS effects on tactile acuity in HCs within the expected plasticity window accounting for task order. Results from ART-ANOVA on 2-PDT task with factors Time (pre, post), ISI(ISI-20, ISI-100), and Order (VTSC-first, 2PDT-first).

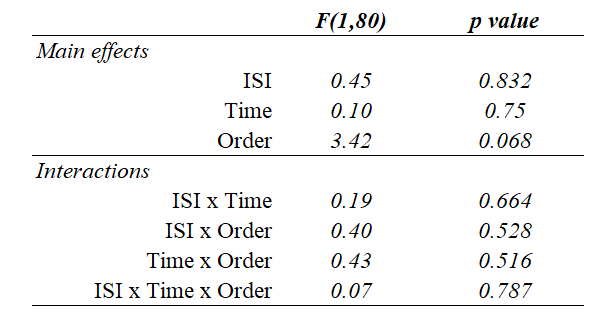
